# Effectiveness of Genomic Selection by Response to Selection for Winter Wheat Variety Improvement

**DOI:** 10.1101/537167

**Authors:** Xiaowei Hu, Brett F. Carver, Carol Powers, Liuling Yan, Lan Zhu, Charles Chen

## Abstract

The genomic revolution opened up the possibility for predicting un-tested phenotypes in schemes commonly referred as genomic selection (GS). Considering the practicality of applying GS in the line development stage of a hard red winter (HRW) wheat variety development program (VDP), effectiveness of GS was evaluated by prediction accuracy, as well as by the response to selection across field seasons that demonstrated challenges for crop improvement under significant climate variability. Important breeding targets for HRW wheat improvement in the southern Great Plains of USA, including Grain Yield, Kernel Weight, Wheat Protein content, and Sodium Dodecyl Sulfate (SDS) Sedimentation Volume as a rapid test for predicting bread-making quality, were used to estimate GS’s effectiveness across harvest years from 2014 (drought) to 2016 (normal). In general, nonparametric algorithms RKHS and RF produced higher accuracies in both same-year/environment cross validations and cross-year/environment predictions, for the purpose of line selection in this bi-parental doubled haploid (DH) population. Further, the stability of GS performance was greatest for SDS Sedimentation Volume but least for Wheat Protein content. To ensure long-term genetic gain, our study on selection response suggested that across this sample of environmental variability, and though there are cases where phenotypic selection (PS) might be still preferential, training conducted under drought stress or in suboptimal conditions could still provide an encouraging prediction outcome, when selection decisions were made in normal conditions. However, it is not advisable to use training information collected from a normal field season to predict trait performance under drought conditions. Further, the superiority of response to selection was most evident if the training population can be optimized.

**Core Ideas:**

- Prediction performance for winter wheat grain yield and end-use quality traits.
- Prediction accuracy evaluated by cross validations significantly overestimated.
- Non-parametric algorithms outperform, when considering cross-year predictions.
- Strategically designing training population improves response to selection.
- Response to selection varied across growing seasons/environments.

## Introduction

Wheat breeding has progressed dramatically in the last century due to the combination of various technologies (Poland et al., 2012); taken together these advancements have driven the yearly genetic gain through selective breeding to nearly a linear increase of 1% in the potential grain yield (Bassi et al., 2016). Faced against human population growth and uncertain climates, global wheat production, however, still falls short (Curtis and Halford, 2014), as the global demand for wheat is projected to increase 60% when the population reaches 9.8 billion by 2050 (Alexandratos and Bruinsma, 2012). The emphasis now is increasingly on not only meeting the food and nutrition demand, but also on how to maximize the opportunity to achieve long-term geo-environmental sustainability responding to local climate challenges.

Genetic improvement has been a major contributor to productivity gains in wheat and other cereal crops (Kharabian-Masouleh et al., 2011). The continued advancement of high-throughput genotyping technologies has stimulated interest in genomic estimated breeding value (GEBV) through genomic selection (GS) (Meuwissen et al., 2001). The use of GEBV for un-typed individuals allows selection to be made solely based on genotypic information; consequently, genetic gain can be largely improved owing to the reduced time and the institutional investment required to complete breeding cycles (Heffiner et al., 2010; Hickey et al., 2014). A large number of GS studies have been conducted, mostly on evaluating the performance of GS algorithms (Desta and Ortiz, 2014; Crossa, et al., 2017), while relatively few considered the schemes of the designated breeding programs (Schulz-Streeck et al., 2012; Zhao, et al., 2012; Gaynor et al., 2017). Summarized from the current knowledge, benefits of adopting GS are determined mainly by the accuracy of estimated GEBV of the validation population (VP) (Hickey et al., 2014; Michel et al., 2017a); when accurate GEBV become available, immediate impact on selecting elites solely by GEBVcould provide a marked decrease in breeding cycle time (Heffner et al., 2009), given that other key breeding targets unamenable to GS are not the limiting factor. However, shortening breeding cycles would lead to a rapid change of genetic diversity in the breeding populations and would affect GEBV accuracy for the long run. Also as pointed out in Bassi et al. (2016) and Crossa, et al. (2017), accurate GS applied in early generations and the opportunities to re-training in later generations determine the long-term, cost-effectiveness of GS outcomes.

The basic form for breeders to predict changes in breeding outcomes is captured in the breeder’s equation (Lush 1937), which describes the evolutionary change in a phenotypic trait by a simple interplay of selection intensity and multi-locus inheritance (narrow sense heritability, *h*^*2*^) (Kelly 2011). This univariate breeder’s equation also states that this expected genetic gain, expressed as the response to selection, is a function of selection intensity, selection accuracy and the standard deviation of breeding values (Falconer and Mackay, 1996). Accuracy of GS has been intensely investigated; however, while low GEBV accuracy might be concerning, response to selection at low GS accuracy is still less variable than response at high accuracy (Hill, 1974; Heffner et al., 2010). The significance of adopting GS can thus be more evident when response to selection per generation is calculated (Resende et al., 2017).

Considering the practicality of GS for wheat improvement programs, emphasis may shift to selecting the elite lines whose superiority in trait performance can be realized in the target environment. At this advanced stage, breeding targets would not be only focus on grain yield and disease resistance, but also include numerous end-use quality traits that are normally laborious and expensive to phenotype (Guzmán et al., 2016; He et al., 2016 and Michel et al., 2017a). Amid sporadic global shortages of high-protein wheat, or more importantly an industry perception of domestic shortages of bread wheat with acceptable dough strength, intensified demand in commercial markets has driven breeding objectives from yield production to end-use quality traits. Hard red winter (HRW) wheat, grown throughout the Great Plains and parts of the U.S. Pacific Northwest and milled into flour for bread, is the biggest U.S. wheat class, typically representing about 40 percent of total U.S. wheat production. Further, human-induced climate change is causing rapid variations of global temperatures and extreme fluctuations in precipitation (Fitzpatrick and Edelsparre, 2018). These changes impose greater pressure for both the crops and breeders, forcing organisms to adapt and breeders to deviate from the established optimal. As a result of the worsening drought in the region, in June 2018, USDA estimated the HRW wheat harvest at 650 million bushels, the second lowest production since 1964. In order to stay competitive in this new wheat market, this study is aimed to examine the effectiveness of GS for both grain yield and protein content phenotypes, using a HRW doubled-haploid (DH) population developed for the southern Great Plains. Factors discussed for GS performance include predictive algorithms and composition of the training population (TP), encouraged by the success in improved predictability when TP is optimized (Rincent et al., 2012; Akdemir et al., 2015; Isidro et al., 2015; Michel et al., 2017a; Neyhart et al., 2017). We evaluated the effectiveness of GS as the response to selection for a more breeder-intelligible assessment of GS performance. Finally, to derive realistic estimates, the response to selection and GS accuracy were assessed with respect to annual precipitation during the growing season that reflects the environmental difference between training and validation populations.

## Materials and Methods

### Duster x Billings Doubled Haploid Population, Field Information and Phenotypes

Developed cooperatively by the Oklahoma Agriculture Experiment Station (OAES) and the USDA-ARS, Duster and Billings are two leading winter wheat cultivars that were released in the Southern Great Plains (Edwards, et al., 2012; Hunger, et al., 2012). A DH population of 282 lines was developed by intercrossing Duster and Billings; agronomic trait performance was assessed in Stillwater, OK, USA (36.12N, 97.09W). The soil type in Stillwater location was Kirkland silt loam or Norge loam (for details, please see http://oaes.okstate.edu/frsu/agronomy-research-station/Stillwater_soilmap.pdf).

The 2014 trial was planted on 11^th^ of November 2013 and harvested on 20^th^ of June 2014. The 2015 trial was planted on 14^th^ of November 2014 and harvested on 14^th^ of June 2015. As for 2016 trial, the planting date was 13^th^ of November 2015 and harvested on 9^th^ of June 2016. The total rainfall between 1^st^ of November and 31^st^ of May was 19.8 cm for the growing season year 2014, 41.3 cm for year 2015, and 45.2 cm for year 2016. In this study, field performance in 2014 trial was referred as drought condition, and 2015 and 2016 as normal. No trials received supplemental irrigation.

In this study, three commonly used phenotypes in the wheat industry, in addition to grain yield (Grain Yield) in kilograms per hectare (kg/ha), were considered; sodium dodecyl sulfate (SDS) Sedimentation Volume (Lorenzo and Kronstad, 1987) adjusted for flour protein (as is) content (SDS Sedimentation Volume), kernel weight measured by the single kernel characterization system (Perten Instruments, Segeltorp, Sweden) (Kernel Weight), and wheat protein on a 12% moisture basis (Wheat Protein). After filtering out missing phenotypic data, 239 DH lines remained for all three years and all four traits; also, only the averaged values from two field replicates of each DH line per year were used in the GS analysis.

### Genotyping-by-sequencing and Genotypic Data Analysis

Three 96-plex libraries were generated from a single-plant sample of each DH line, with three replicates of each parents. A library consisting of DNA fragments with a forward adapter and a reverse adapter on opposite ends of every fragment was produced based on the protocols of Poland et al. (2012a) using the combination of *Pst*I (CTGCAG) and *Msp*I (CCGG) restriction enzymes to perform genome complexity reduction. The fall list of barcoded adapters and the corresponding DH samples can be found in Li et al. (2015). Detailed procedures of library construction were described in Poland et al. (2012a).

In brief, PCR products were amplified using short extension time (< 30 seconds) to enrich shorter fragments suitable for bridge-amplication on the Illumina flowcell on Illumina HiSeq2000. The raw reads were assigned to individual samples based on an exact match to the DNA barcode followed by the *PstI* restriction site, and trimmed to 64 base-pairs. The tag sequences were aligned allowing a one or two base-pair mismatch for SNP determination. When a SNP call showed a difference between two alleles, a genotyping-by-sequencing (GBS) SNP was considered. Due to the nature of DH lines, when the SNP call was identified as heterozygous, information about the locus was retained but the genotypic data was set to missing. In total, 14,028 SNPs were generated from this SNP determination procedure.

GBS is a genotyping technology with high density but with potentially low genome coverage as well. On average, the mean missing ratio is 42% per SNP locus per individual, with the maximum missing ratio as high as 78%. To obtain quality genetic markers for prediction, SNP markers were filtered based on: 1) less than 50% missing ratio, and 2) at least 5% minor allele frequency (MAF), totaling 7,426 SNPs for further analysis. Rutkoski et al. (2013) showed that relative reliable performance on wheat genotypes using the map-independent imputation method, like k-nearest neighbor (kNN) and random forest (RF). Following SNP determination, kNN was used to impute missing SNP data in this study owing to its computational efficiency; the kNN imputation used in this study was implemented in R package *scrime* (Schwender 2013).

### Genomic Selection Algorithms

Genomic selection exploits a statistical model which partitions phenotypic variation into genetic and non-genetic components. In the model, the phenotypic value of an individual is assumed to follow a Gaussian distribution with the center at the linear function of its genotypic values. The statistical model is shown in equation (1).

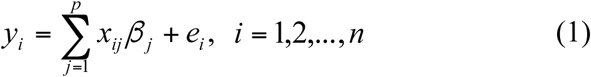

where *y*_*i*_ is the standardized phenotypic value of the *i*th individual so that the intercept term is omitted for all traits; *x*_*ij*_ encodes the genotype of the *i*th individual at marker locus *j* = 1,…, *p.* In *x*_*ij*_, the allelic states of individuals are coded as 0 and 2 for diploid genotypic values of AA and aa, respectively; *β* _*j*_ is the marker’s effect at the *j*th locus; and the error term 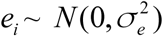. The genomic estimated breeding value of an individual, GEBV, is thus defined as the predicted phenotypic value based on this linear model. With high-density marker data like GBS, the number of markers (*p*) is larger than the number of individuals or records (*n*). Owing to this “*small n, large p*” conundrum, statistical methods that apply penalty or regularization through constraints in the objective function by variable selection, or by introducing prior distributions for unknown parameters in a Bayesian framework, are broadly adopted in GS algorithms for estimating marker effects. In this study, GS algorithms of choice were Bayesian LASSO (BL) (Park and Casella, 2008), Random Forest (RF) (Breiman 2001), Reproducing Kernel Hilbert Space (RKHS) (Gianola et al., 2006), and Ridge Regression Best Linear Unbiased Prediction (RRBLUP) (Hoerl and Kennard, 1970).

### BL (Bayesian LASSO)

BL utilizes a conditional mixture of Gaussian distributions that describes the prior distribution of the marker effect. Consequently, the assumption of equal variance across all markers is relaxed. The marker-specific priors are modeled by the following hierarchical distributions.

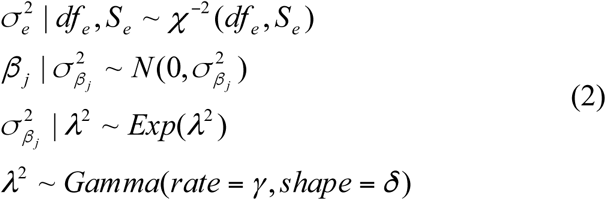

In (2), a large *λ*^2^ value will lead to a sharp prior distribution for *β* _*j*_ centered at 0 through a reduced variance, therefore, more shrinkage towards zero. The effect due to the choice of *λ*^2^ in the BL has been addressed in de los Campos et al. (2009a) and Lehermeier et al. (2013). These authors studied the influence of the choice of hyperparameters in the Gamma distribution for *λ*^2^ and concluded that, even the influence of *λ*^2^ is unknown, the model goodness-of-fit and the estimates of genetic values were quite robust with respect to the choice of *λ*^2^ R package *BGLR* (de los Campos and Pérez, 2013) was used for BL model fitting with 100,000 iterations in total, 30,000 iterations for burn-in, and 50 for thinning. The convergence of the Markov chains was confirmed by visualizing the sampling paths.

### RF (Random Forest)

An RF predictor is an ensemble of individual classification or regression tree predictions (Breiman 2001). Each individual tree was grown on bootstrap samples of observations using a random subset of predictors to define the best split at each node. The RF prediction for an observation is computed by averaging the predictions over trees for which the given observation was not used to build the tree (Heslot et al., 2012). This algorithm was implemented in R package “RandomForest” (Liaw and Wiener, 2002).

### RKHS (Reproducing Kernel Hilbert Space)

RKHS uses a kernel function to convert SNP marker data into a set of genetic distances between pairs of observations, resulting in a square matrix that can be used in a linear model. An advantage of RKHS is that no linearity is assumed; therefore, in theory, non-additive effects can be better captured. The Bayesian RKHS regression regards genetic values as random variables coming from a Gaussian process centered at zero and with a covariance structure that is proportional to a kernel matrix *K* (de los Campos et al., 2010); that is, *Cov*(*g*_*i*_, *g* _*j*_) ∝ *K* (*x*_*i*_, *x* _*j*_), where *g*_*i*_, *g* _*j*_ are the sum of genetic values for the *i*th and *j*th individuals, respectively; *x*_*i*_, *x* _*j*_ are vectors of marker genotypes for the *i*th and *j*th individuals, respectively, *i*.*e*., *x*_*i*_ = (*x*_*i*1_,…,*x*_*ip*_)′ and *x* _*j*_ = (*x* _*j*1_,…,*x* _*jp*_)’; *K* (.,.) is a positive definite function evaluated in marker genotypes. The kernel function used in this study was adopted from de los Campos et al. (2010),

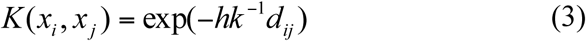

where *h* is a bandwidth parameter that controls the rate of decay of the correlation between genotypes; *d*_*ij*_ = ||*x*_*i*_ – *x* _*j*_||^2^ is a marker-based squared Euclidean distance between two individuals *i* and *j*; and 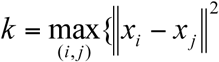. The optimal value of *h* that produces the maximum prediction accuracy was selected by grid search over 5-fold cross-validation (CV). This algorithm was implemented in R package *BGLR* (de los Campos and Pérez, 2013). In this study, the predictability of Bayesian RKHS was tested using 12,000 iterations, and 2,000 for burn-in. The convergence of the Markov chains was confirmed by visualizing the sampling paths.

### RRBLUP (Ridge Regression Best Linear Unbiased Prediction)

In penalized regression algorithms, a penalty function is used to constraint the size of the estimated regression coefficients in the objective function. The estimated marker effects can be obtained as

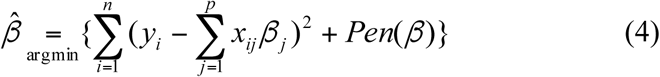

The penalty function of RR is defined by *L*_2_ norm of the regression coefficients, *i*.*e*., 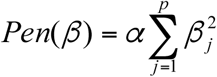 where *α* is the penalty parameter. So the estimated marker effects can be obtained as

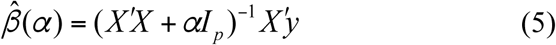

where *X* is *n* × *p* design matrix of SNP marker data.

This assures a unique inverse and stabilizes the solution when *X′X* is ill-conditioned due to multi-collinearity (Hoerl and Kennard, 1970). The bias of the estimated regression coefficients increases but the variance decreases monotonically with *α,* and mean squared error (MSE) can be improved compared to the least squares estimation (Hoerl and Kennard, 1970). RRBLUP performs no variable selection and thus retains all markers in the model. The choice of appropriate penalty parameter *α* affects the performance of penalized regression methods. The optimal value of *α* that minimizes MSE was selected by 5-fold CV using the training samples. This algorithm was implemented in R package *glmnet* (Friedman et al., 2009).

### Genomic Selection Algorithm Evaluation

The prediction performance of GS algorithms was evaluated by using 5-fold CV. Each phenotypic dataset was randomly divided into 5 equal parts. Then the GEBVs for each fold were predicted by training the model on the four remaining folds. The entire procedure was repeated 5 times, such that GEBVs of all individuals can be obtained. Pearson’s correlation coefficient (PCOR) and MSE between observed phenotypic value and GEBV were used to assess the performance of prediction algorithms.

### Response to Selection

In the line-testing stage of HRW wheat variety development program (VDP), no new genetic diversity is introduced and selection pressure shifts from populations to experimental lines. The focus of such programs will be on a stable selection of superior lines over years. For that reason, we introduced a procedure that evaluates line selection by relative superiority of selection, as a measurement of response to selection (Michel et al., 2018). Suppose *S* top performers (*i*.*e*., lines with desirable values, high or low) to be selected, the relative superiority of selection, 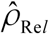, is defined as below (Michel et al. (2018)):

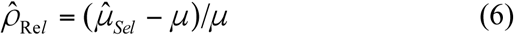

where *μ* is the average phenotypes of the entire population; 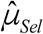 is the estimated average phenotypes of the selected population. And by Michel et al. (2018),

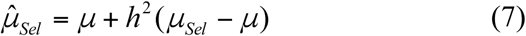

where *h*^2^ is the heritability of the trait; *μ*_*Sel*_ is the average phenotypes of the selected population. The heritability *h*^2^ was set to 1 when evaluating GS; the estimated heritability from a linear mixed model (detailed below) was used when phenotypic selection (PS) was considered. As a result, the performance of selected lines can then be evaluated by changes in 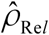 over growing seasons in the line testing stage of winter wheat breeding programs. Responses to selection due to selection intensity were explored by selecting three different proportions of top performers from the entire DH population, denoted by *S* (*i*.*e*., *S* =10%, 30%, and 50%).

The heritability of each phenotypic trait was estimated from the following linear mixed model.

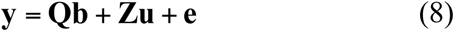

where **y** is a *qn* ×1 vector of *q* years’ phenotypes; **Q** is a *qn* × *q* design matrix of the fixed year effect, **b** is a vector of the fixed year effect; **Z** is a *qn* × *n* design matrix of random genetic effects, a vector of random genetic effect 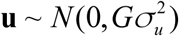, the genomic relationship matrix *G* is calculated by a linear kernel matrix 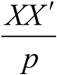 where *X* is an *n* × *p* design matrix of SNP marker data; the error term 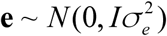. Then the narrow-sense heritability *h*^2^ was estimated as 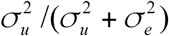. The estimation was implemented in R package *sommer* (Covarrubias-Pazaran 2016).

### Design of Training Populations

To further understand the effectiveness of GS in the line testing stage, the capacity for customizing the training information was examined. The principle applied in this research objective was to maximize the likelihood of capturing the underlying QTLs (quantitative trait loci). To do so, we refer to this as design of training population (DTP), a procedure to capture as much phenotypic variance as possible. Suppose one wishes to compose a DTP with only *m* lines from the entire population such that a smaller DTP can be constructed in order to accommodate more field sites; the selection of DTP is then made of (1 / 2)*m* lines with the highest values, and another half with the lowest values.

For the trait of interest, three phenotypes per DH line could be used to construct the DTP: observed phenotype, GEBV, and a hybrid method of using both phenotypes and GEBVs; these were termed “DTP_Ptails”, “DTP_Gtails” and “DTP_Htails”, respectively. “DTP_Ptails” selects (1 / 2)*m* lines with the highest and lowest values respectively based on the observed phenotypes to form DTP; “DTP_Gtails” uses GEBVs, instead of observed phenotypes; and, “DTP_Htails” uses the majority votes from the “DTP_Ptails” and “DTP_Gtails”. In this study, four different scenarios of *m* (*i*.*e*., *m* = 20%, 40%, 60%, and 80% of the entire population) were explored for the training population (TP) size. For proof of concept, only the results from the best GS algorithm were used to obtain average GEBVs for each trait. Finally, the DTP were used to train the model for updated GEBVs, and then these updated GEBVs were used for selection. The performance of DTP was evaluated by the relative superiority of selection 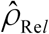 over growing seasons. Finally, to show the effect of reduced TP on response to selection, we also compared the performance of GS by DTP versus GS by all training population (GS_ATP).

## Results

### Summary of Phenotypic Data and Heritability

The empirical correlation of phenotypes between years of all traits can be found in Table 1. Overall, SDS Sedimentation Volume had the highest empirical correlation between years averaged across pairs of years, 0.58 (0.52 – 0.70), while Wheat Protein owned the lowest (0.17-0.36, average of 0.26). Additionally, the phenotype average and standard deviation (SD) across years are presented in Table 2 for all traits. Grain Yield showed the largest variation among years, while SDS Sedimentation Volume was relatively stable over years.

**Table 1.** Raw phenotypic correlation between years for four traits

| Trial Year | **Grain Yield** |  |  | **SDS Sedimentation Volume** |  |  |
| --- | --- | --- | --- | --- | --- | --- |
|  | 2014 | 2015 | 2016 | 2014 | 2015 | 2016 |
| 2014 | 1 | 0.42 | 0.24 | 1 | 0.53 | 0.52 |
| 2015 |  | 1 | 0.43 |  | 1 | 0.7 |
| 2016 |  |  | 1 |  |  | 1 |
| Trial Year | **Kernel Weight** |  |  | **Wheat Protein** |  |  |
|  | 2014 | 2015 | 2016 | 2014 | 2015 | 2016 |
| 2014 | 1 | 0.26 | 0.4 | 1 | 0.17 | 0.36 |
| 2015 |  | 1 | 0.44 |  | 1 | 0.25 |
| 2016 |  |  | 1 |  |  | 1 |

**Table 2.** Descriptive summary and estimated heritability of four phenotypic traits tested

| Trial Year | Mean (SD) | Three-year $\hat{h}^2$ | Mean (SD) | Three-year $\hat{h}^2$ |
| --- | --- | --- | --- | --- |
| **Grain Yield** |  |  | **SDS Sedimentation Volume** |  |
| 2014 | 1,577.50 (270.17) | 0.52 | 7.18 (0.55) | 0.70 |
| 2015 | 2,209.44 (441.28) |  | 7.73 (0.73) |  |
| 2016 | 2,471.35 (720.55) |  | 7.41 (0.74) |  |
| **Kernel Weight** |  |  | **Wheat Protein** |  |
| 2014 | 25.48 (1.58) | 0.67 | 13.48 (0.71) | 0.54 |
| 2015 | 24.36 (2.16) |  | 12.32 (0.51) |  |
| 2016 | 31.80 (2.63) |  | 12.08 (0.58) |  |

Table 2 shows the estimated trait heritability obtained from the linear mixed model with three years’ phenotype information. SDS Sedimentation Volume was highly heritable (0.74), while Grain Yield and Wheat Protein were moderately heritable (0.52 and 0.54, respectively).

### Performance Assessment of GS Algorithms

To validate GS prediction the average PCOR and MSE, as well as their SD, were generated from 100 replications of 5-fold CV. Results of prediction accuracy are shown in Figure 1 and Figure 2 (for PCOR); the variability of these results are included in the Supplementary Fig. S5 – S6 (for MSE). Overall, the GS algorithm with the highest PCOR correlation tended to show the lowest MSE. Our BL and Bayesian RKHS results all reached convergence of Markov chains (Supplementary Fig. S1 – S4).

**Figure 1:**
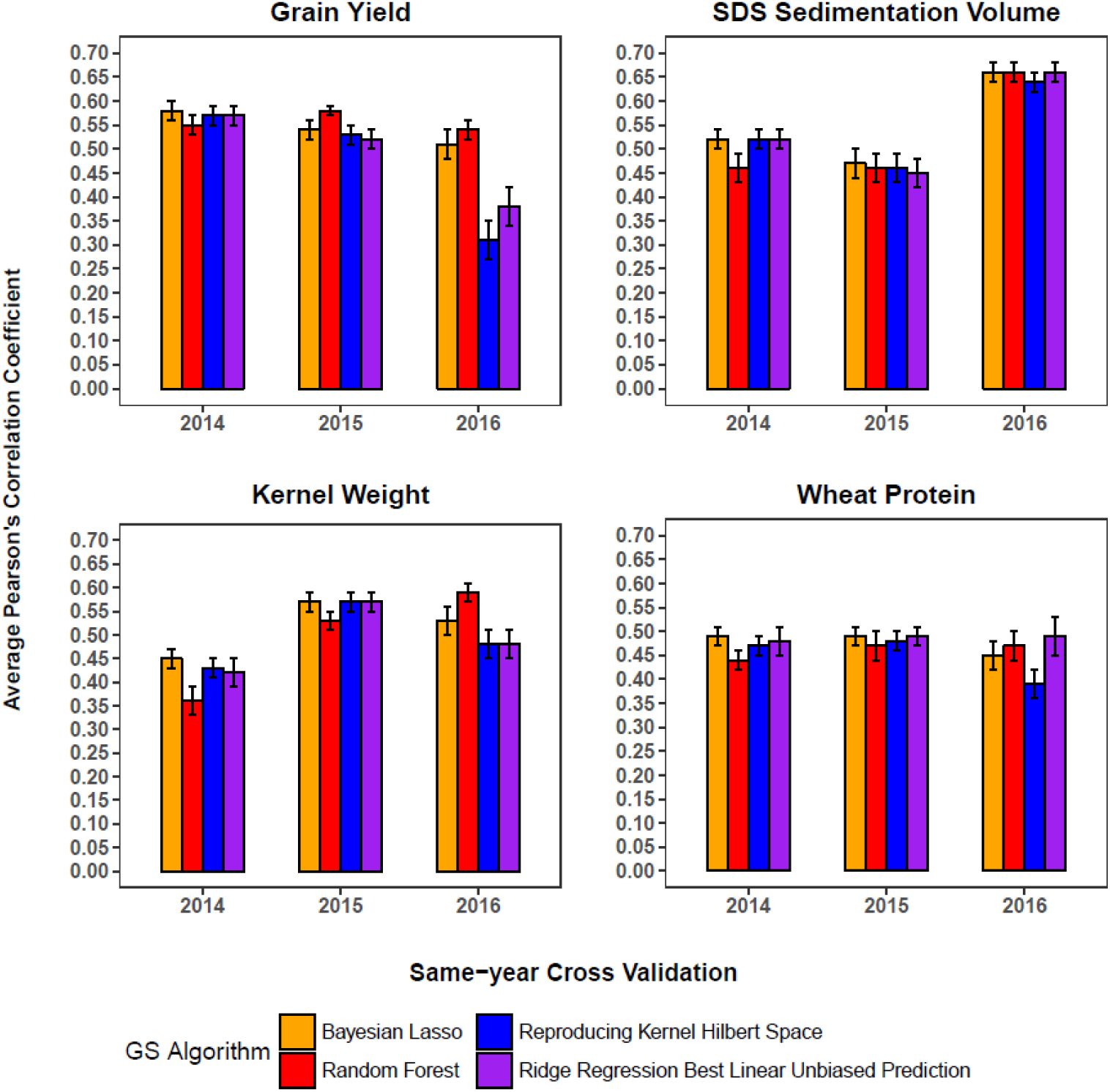
Average Pearson’s correlation coefficient of same-year cross validation for Grain Yield, SDS Sedimentation Volume, Kernel Weight, and Wheat Protein. The error bar stands for one standard deviation.

**Figure 2:**
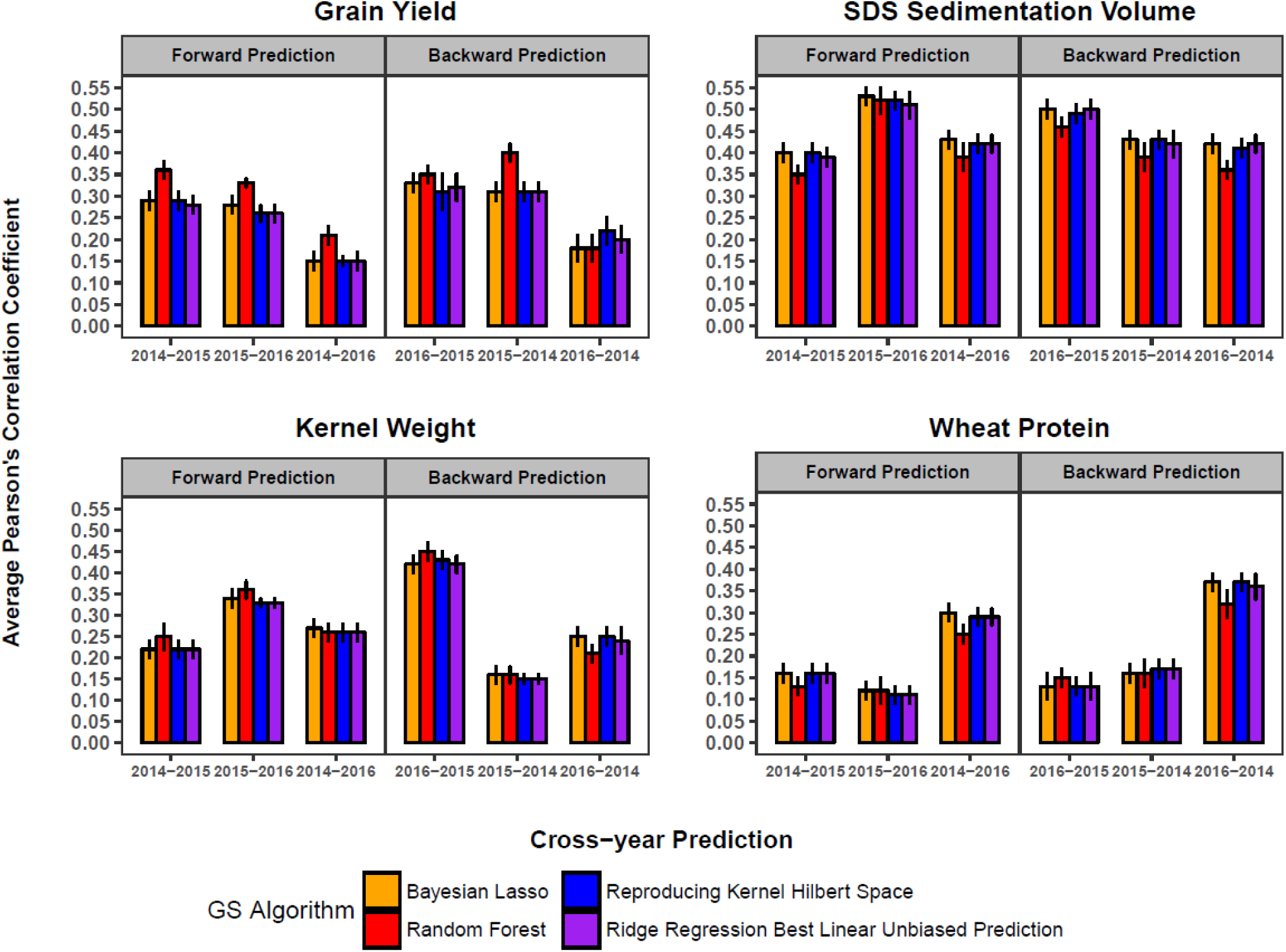
Average Pearson’s correlation coefficient of cross-year prediction for Grain Yield, SDS Sedimentation Volume, Kernel Weight, and Wheat Protein. The error bar stands for one standard deviation. The year before and after dash stands for training and validation year respectively.

### Same-year Cross-validations

For Grain Yield, the average PCOR (MSE) of three years was 0.54 (0.7), 0.56 (0.7), 0.47 (0.76), and 0.49 (0.75) from BL, RF, RKHS, and RRBLUP, respectively. BL and RF performed similarly and resulted in 9% higher predictability than RKHS and RRBLUP. It is however worth noting that the significantly lower prediction accuracies of RKHS and RRBLUP were caused by year 2016. The four algorithms performed similarly in year 2014, whereas in all comparisons RF produced slightly higher accuracies in year 2015.

For SDS Sedimentation Volume, the average PCOR and MSE of three years were similar among four algorithms, with the exception of the significantly lower GS accuracy for RF in year 2014. Overall, the average same-year PCOR was highest in year 2016.

The average PCOR (MSE) of three years were 0.52 (0.73), 0.49 (0.75), 0.49 (0.75), and 0.49 (0.76) from BL, RF, RKHS, and RRBLUP, respectively for Kernel Weight. BL outperformed slightly, when average PCOR was used for comparison. Although the average performances of RF, RKHS, and RRBLUP were found similar, RF results varied significantly among growing seasons; among all GS algorithms, RF arrived to the lowest single year GS accuracy for Kernel Weight at PCOR=0.36 in 2014 but highest PCOR at 0.59 in 2016.

On average, Wheat Protein GS prediction was at 0.47, with the lowest GS found in 2016 using RKHS (PCOR=0.39). GS algorithm performance was consistent between MSE and PCOR, whereas BL and RRBLUP showed slightly lower average MSE at 0.77, and it was slightly higher for RF and RKHS (MSE=0.79, Fig. S5).

In summary, average prediction accuracies across four GS algorithms and three years was highest for SDS Sedimentation Volume (0.54) and lowest for Kernel Weight (0.47).

### Cross-year Predictions

To examine applicability of GS in a more realistic setting, GS performance was trained in one growing season, and then examined in another season; this can also be interpreted as GS predictability across environments. Also, rainfall is an unpredictable weather factor in arid areas like the southern Great Plains of the US. Since field management of the research station site remained consistent among years, and there was no new genetic diversity introduced, the main driving factor for the line performance, we reckoned, was variability in total precipitation among growing seasons.

For clarification, we split the results into two groups, forward prediction and backward prediction. Forward prediction was used to describe the scenarios where the previous year was used as training populations (TP) to predict future years. For example, the prediction of year 2015 field evaluation, GS was applied using 2014 as training; this scenario was denoted as “2014–2015” in Figure 2. Backward prediction is reverse. Although forward predictions are consistent with practical application, backward predictions were examined to consider GS applicability of cross-environmental variability. Figure 2 and Supplementary Fig. S6 showed PCOR and MSE, respectively, for cross-year predictions.

In forward predictions (*e*.*g*., 2014 – 2015, 2015 – 2016, 2014 – 2016), the average PCORs for grain yield over these comparisons were 0.24, 0.30, 0.23, and 0.23 from BL, RF, RKHS, and RRBLUP, respectively (Figure 2); the corresponding average MSEs were 1.07, 0.99, 1.03, and 1.05 (Fig. S6). Similar results of average PCORs and corresponding MSEs were observed in backward predictions (*e*.*g*., 2016 – 2015, 2015 – 2014, 2016 – 2014). Overall, RF performed significantly better among the GS algorithms tested, regardless of prediction scenario.

In all cross-year predictions, SDS Sedimentation Volume generated the most consistent GS results across growing seasons and across GS algorithms. Performance evaluations with average PCOR and MSE were similar for all algorithms, except for RF (Figure 2). However, GS performance for RF was only significantly lower in 2014 – 2015 and 2016 – 2014 (∼ 0.36, whereas others averaged 0.45). RF also exhibited the highest MSE, though not significant.

Similar to SDS Sedimentation Volume, there was no observable performance difference in GS algorithms for Kernel Weight; the average PCOR for Kernel Weight were comparable across prediction scenarios (Figure 2). However, GS prediction accuracy of 2016 2015 was significantly greater than other comparisons (∼0.45); and, the 2015 2014 backward prediction produced a low accuracy 0.16.

For Wheat Protein, all GS algorithms performed closely in average accuracy for both forward and backward predictions, except that RF showed a slight disadvantage in 2016 – 2014. The best GS algorithm for Wheat Protein is either BL or RKHS; however, RKHS showed a minor advantage in MSE (Fig. S6). Similar with Grain Yield, backward prediction accuracies were greater than forward predictions for Wheat Protein. With the best predictable scenario, the average PCOR for Wheat Protein was at 0.37 for 2016 – 2014.

### Computational Efficiency

The differences in computing time among these four GS algorithms in our study were considerable. For example, to finish both same-year and cross-year GS predictions over 100 replications of 5-fold CV, 1.4 hours were required for RRBLUP, 1.81 hours for RKHS, 3.19 hours for RF; BL was the most computationally intense, requiring 270 hours for completion.

### Relative Superiority of Selection

Based on our evaluation in same-year and cross-year GS performances, RF was used to study changes in response to selection over the growing seasons for Grain Yield and Kernel Weight; RKHS was selected for SDS Sedimentation Volume and Wheat Protein. For each trait, changes on relative superiority of selection 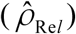, with respect to size of the training population constructed from the GEBVs of DTP (GS_DTP_Gtails; purple lines in the Figure 3–6) and the majority votes using both of the GEBVs and observed phenotypes (GS_DTP_ Htails; green lines in the Figure 3–6), were evaluated relative to the use of the entire training population (GS_ATP; blue lines in the Figure 3–6). Since there is no optimization or reduction of training for GS_ATP and PS (red lines in the Figure 3–6), relative superiority of selection from these two selection methods remained unchanged across *m* values (Figure 3–6).

**Figure 3:**
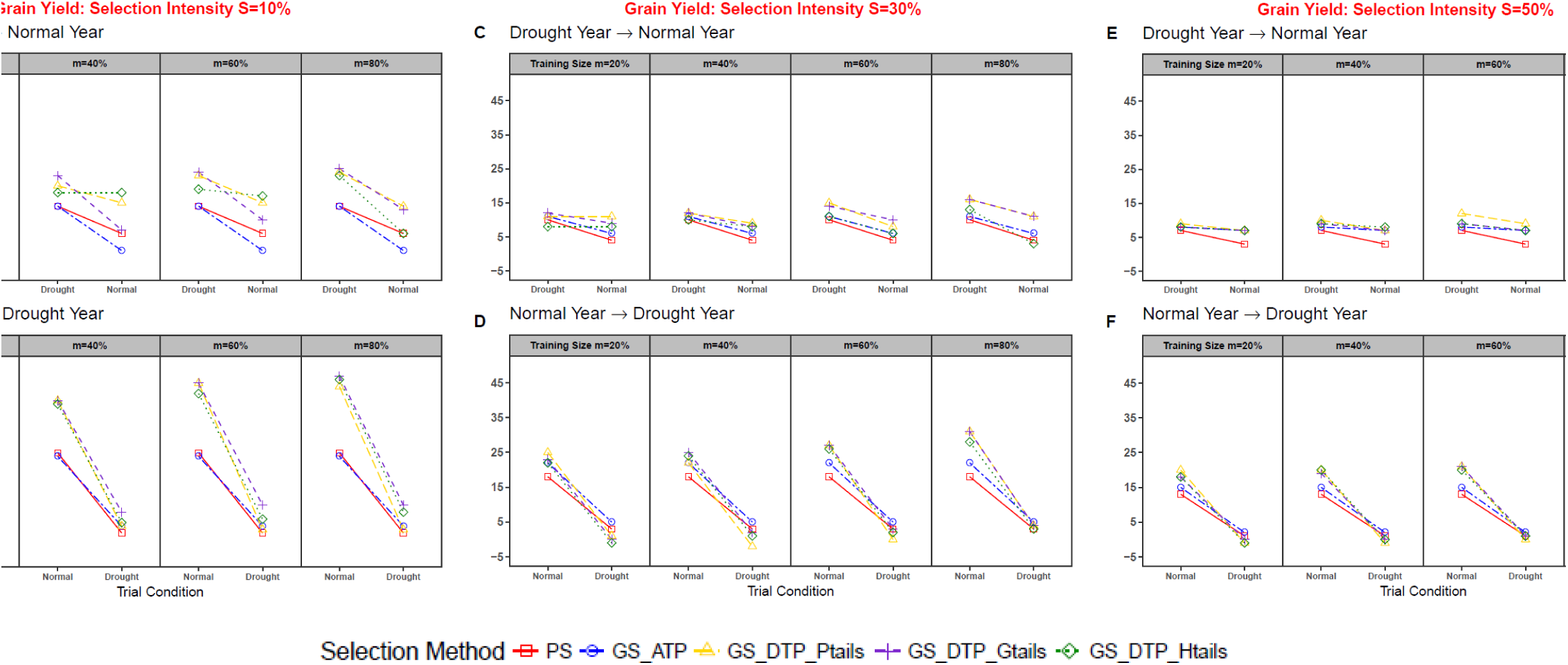
Relative superiority of selection for trait Grain Yield. PS, phenotypic selection; GS_ATP, genomic selection by all training population; GS_DTP_Ptails, genomic selection by designed training population with phenotypic tails; GS_DTP_Gtails, genomic selection by designed training population with genomic tails; GS_DTP_Ptails, genomic selection by designed training population with hybrid tails.

**Figure 6:**
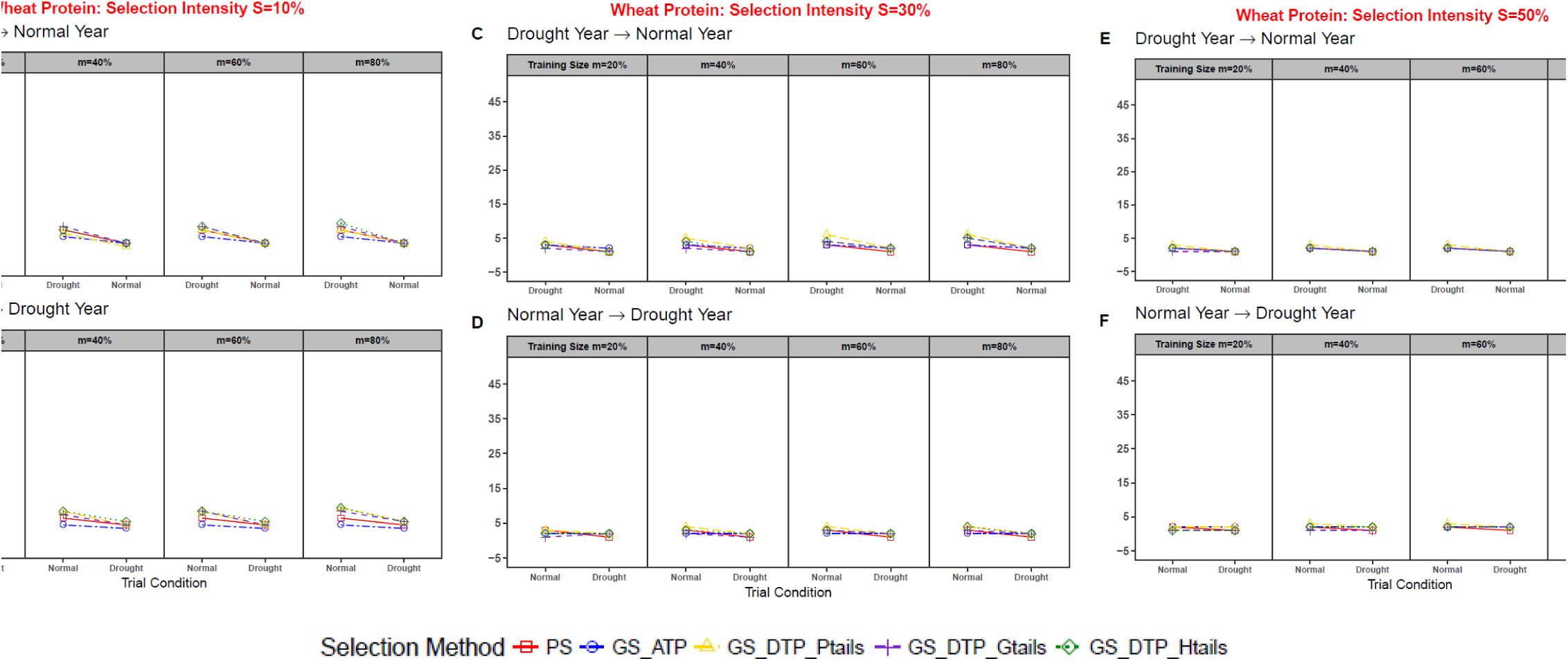
Relative superiority of selection for trait Wheat Protein. PS, phenotypic selection; GS_ATP, genomic selection by all training population; GS_DTP_Ptails, genomic selection by designed training population with phenotypic tails; GS_DTP_Gtails, genomic selection by designed training population with genomic tails; GS_DTP_Ptails, genomic selection by designed training population with hybrid tails.

To investigate effects of environmental variability to the response of selection for GS, changes in 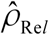 over growing seasons were evaluated in two scenarios, the superior line was selected initially in a drier condition/year (*i*.*e*., year 2014) and then evaluated in a well-irrigated growing condition/year (*i*.*e*., year 2016) (Drought year **→** Normal year in the Figure 3–6); and the reverse order (Normal year **→** Drought year in the Figure 3–6). Overall, a gradual reduction in response to selection was observed in all scenarios, as response to selection was greatest at the initial selection year (or growing environment) and completely diminished in some cases at the final testing year. Also, note that the red lines in the Figure 3–6 denote the response to selection done only with phenotypic information, a scenario replicating current conventional breeding practice. Effectiveness of adopting GS was therefore examined by comparing changes in the response to selection 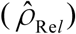 to the scenarios represented by the corresponding red lines. A number of scenarios where GS might not be desirable was also documented (*e*.*g*., GS_ATP in Figure 3A). Additionally, no noticeable difference was detected between GS and PS in some scenarios (*e*.*g*., Figure 6).

#### Grain Yield

As expected, the greatest GS performance relative to selection response was found at the initial selection, when selection was conducted non-drought growing conditions and when a large training population is available (*m* = 80%, Normal **→** Drought Year, Figure 3B, 3D, and 3F); and as the target growing condition for validation population began to deviate from that of the training population, relative response to selection was reduced; this is common for all selection scenarios for Grain Yield (Figure 3). Under the same scenario (Normal **→** Drought Year), the benefit of adopting GS can also be seen in cases where greater selection intensity was applied (Figure 3B); a 47% and 46% relative superiority of response to selection resulted when applying GS directly (GS_DTP_Gtails) and in combination with phenotypic selection (GS_DTP_Htails), respectively, compared to the 20% response to selection of *S* = 50% (Figure 3F).

Considering the growing condition used for training for GS, reduced performance of GS was observed for predicting Grain Yield with information trained in the drought-stressed environment (Drought **→** Normal Year, Figure 3A, 3C, and 3E). Using models trained under this condition, GS performance in terms of selection response was hampered; only 25% of 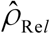 could be obtained as the best scenario with a greater selection intensity (*S* = 10%) and a large training population (*m* = 80%), as exhibited in Figure 3A. When selection intensity was lessened (*S* = 30%), a larger training population was required to obtain 13 – 16% of relative superiority of selection response with only GS data used (GS_DTP_Gtails in Figure 3C); further lessening the selection intensity would further diminish the advantage for using GS, and difference in the selection response between phenotypic selection and GS was reduced to as little as 1 – 3% (Figure 3E). Amongst all scenarios in Drought **→** Normal Year, it is worth noting that, when selection intensity was high (*S* = 10%), the benefit of using GS was not suggested in the case of GS_ATP, the use of the entire training population without optimizing the genetic architecture underlying for the variation of Grain Yield.

#### SDS Sedimentation Volume

Unlike Grain Yield, the change of 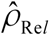 over growing seasons for SDS Sedimentation Volume was relatively stable in comparison; when selection was done in a normal growing season, the average response to selection across growing seasons that experienced drought was 10%, 5%, and 2% respectively for *S* = 10%, 30%, and 50% (Normal **→** Drought year, Figure 4B, 4D, and 4F). When selection intensity was high (*S* = 10%), a sizable superiority can also be observed in the overall differences in 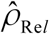 between GS and PS, demonstrating an average of 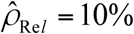 for the final testing year using GS compared to PS alone (Figure 4B). Also, predicting with a larger training population showed a benefit for GS (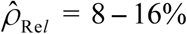 at the initial selection year from *m* = 80%, Figure 4B, 4D, and 4F). When selection was performed in a drought-stressed environment (Drought **→** Normal year, Figure 4A, 4C, and 4E), overall response to selection was insignificant in the cases of low selection intensities (*S* = 30% and 50%, Figure 4C and 4E). Also, under the same Drought **→** Normal year scenario, sizeable training information was required for GS to be effective for higher selection intensity (*S* = 10%, Figure 4A).

**Figure 4:**
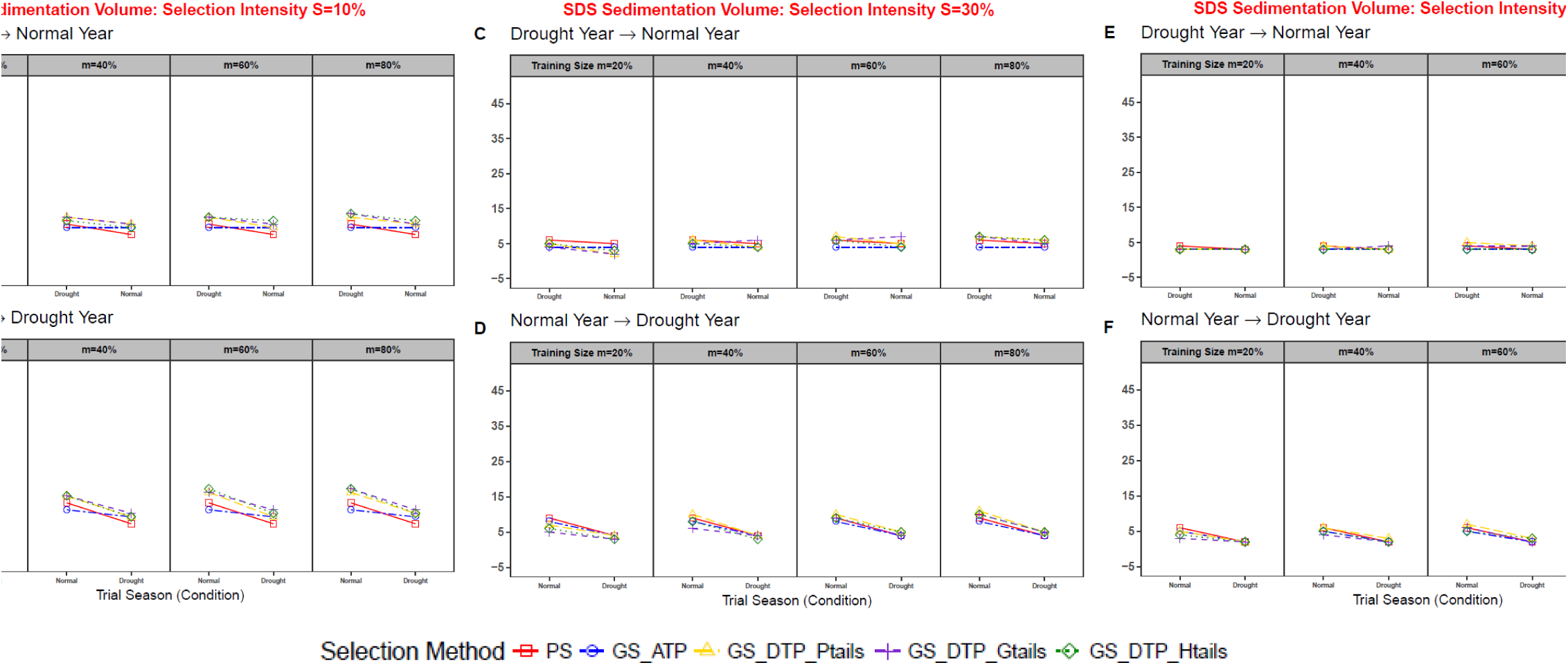
Relative superiority of selection for trait SDS Sedimentation Volume. PS, phenotypic selection; GS_ATP, genomic selection by all training population; GS_DTP_Ptails, genomic selection by designed training population with phenotypic tails; GS_DTP_Gtails, genomic selection by designed training population with genomic tails; GS_DTP_Ptails, genomic selection by designed training population with hybrid tails.

#### Kernel Weight

Shown in Figure 5, evaluation of GS scenarios on Kernel Weight was similar to SDS Sedimentation Volume, *i*.*e*., with lower selection intensities of *S* = 30% and 50%, changes in 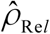 were insignificant for Drought **→** Normal Year (Figure 5C and 5E). Applying a greater selection intensity (*e*.*g*., *S* = 10%), the change in 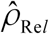 over growing seasons for GS became more evident, when selection was done in normal conditions (Normal **→** Drought Year, Figure 5B). Overall, the relative responses to selection averaged 7%, 5%, and 4% for *S* = 10%, 30%, and 50%, respectively for Normal **→** Drought Year scenarios (Figure 5B, 5D, and 5F). Also, our results showed that an average 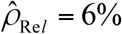 could be expected from GS for Kernel Weight, when a large training population is available (*m* = 80%, Figure 5).

**Figure 5:**
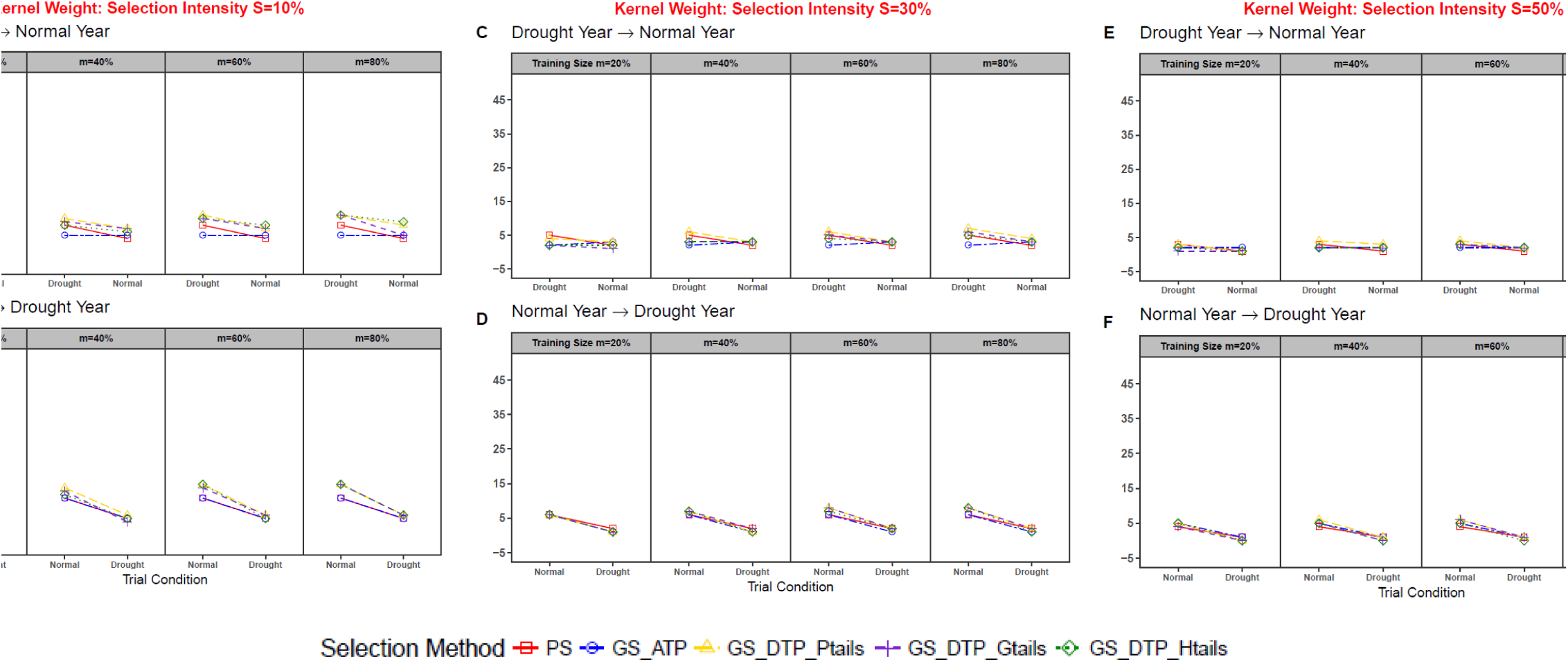
Relative superiority of selection for trait Kernel Weight. PS, phenotypic selection; GS_ATP, genomic selection by all training population; GS_DTP_Ptails, genomic selection by designed training population with phenotypic tails; GS_DTP_Gtails, genomic selection by designed training population with genomic tails; GS_DTP_Ptails, genomic selection by designed training population with hybrid tails.

#### Wheat Protein

Results of relative superiority of selection for Wheat Protein were showed in Figure 6. Overall, GS performance as response to selection for Wheat Protein was insignificant; no marked difference was observed between two selection scenarios and among three selection intensities. Averaged at 2% overall, response to selection was the lowest amongst the four traits examined. Even at the initial selection year, the resulting response to selection was as little as 4% with the greatest selection intensity (Figure 6A and 6B).

## Discussion

Provided with unprecedented genotyping capacity, the emergence of GS has overcome some shortcomings of marker assisted selection (MAS) (Howard et al., 2014), shifting the paradigm of how breeding decisions are made (Meuwissen et al., 2001). Supported by numerous studies (*e.g.* Desta et al., 2014; Beyene et al., 2015; Rutkoski et al., 2015; Battenfield et al., 2016; He et al., 2016; Song et al., 2017; Michel et al., 2018), GS has a promising future in plant breeding (Jonas and de Koning, 2013).

A number of interrelated factors affect GS prediction performance. These include the size of the training population (Rincent et al., 2012; Akdemir et al., 2015), relatedness between training and validation populations (Rincent et al., 2012; Akdemir et al., 2015; Isidro et al., 2015; Michel et al., 2017a; Neyhart et al., 2017), marker density (Daetwyler et al., 2010), heritability (Heffner et al., 2009), genetic architecture of the target trait and the distribution of linkage disequilibrium (LD) between genetic markers and the underlying quantitative trait loci (QTL) (Desta and Ortiz, 2014). In the original paper of GS, Meuwissen et al. (2001) studied numbers of QTLs and relative magnitude of genetic effects and suggested that, without interaction terms, improved accuracy can be achieved if distribution of genetic effects was known. Similarly, including the effect of major genes showed advantages for GS performance in a simulation study; the advantage of GS however begins to diminish when the number of QTLs is greater than 10 (Bernardo 2013). In cases where full-sib families are used, performance of GS algorithm evaluated by cross-validations within the population would be determined by the presence of major genes, or whether or not marker density is enough to capture the LD with QTLs (Howard et al., 2014).

Considering GS accuracy estimated by same-year cross-validations, performance of Grain Yield, a classic polygenic trait, showed a high degree of variability among GS algorithms used. BL and RF demonstrated stable predictability in all same-year cross-validations for Grain Yield (Figure 1), whereas RRBLUP and RKHS showed much greater variation of prediction performance among three years. BL and RRBLUP are both linear models that assume the linearity of marker effects. The difference between these two lies primarily in the shrinkage of marker effects, where RRBLUP assumes equal variance and shrinks all the marker effects to the same level and BL can actually shrink some coefficients exactly to zero, performing as a variable selection method. With a relatively smaller sample size in our study, we suspect that the superior performance of BL and RF was due to their ability to capture the non-uniform distribution of marker effect across the genome (Daetwyler et al., 2010), while avoiding overfitting. This advantage was also evident in cross-year predictions (Figure 2). Using a large European winter wheat population of 2,325 commercial lines in He et al. (2016), the polygenic nature of Grain Yield is more likely to be captured by the sample size; as a result, these associated issues of RRBLUP appeared to be alleviated in He et al. (2016).

As for end-use quality traits like milling and baking quality, indirect phenotyping on correlated traits like protein content has been used as selection criterion (Michel et al., 2018). Most of these traits are considered to be governed by a mixture of major genetic loci and many other small-effect loci (Giroux and Morris, 1998; Olmos et al., 2003; Lillemo et al., 2006; Heffner et al., 2011; El-Feki et al., 2013; Battenfield et al., 2016; Würschum et al., 2016; Michel et al., 2017b). For example, pre-selection of the known *Glu-1* and *Glu-3* glutenin loci in early generations as a marker-assisted approach has shown some merit (Kuchel et al., 2007; Krystkowiak et al., 2016), but the number of successful cases is few (Michel et al., 2018), mainly due to the incapacity to include the genetic variance contributed by a large number of small-effect loci (Reif et al., 2011; Tsilo et al., 2011, 2013; Cabrera et al., 2015), as well as to the complex interaction of allelic effects on dough rheology and bread-making properties (Langner et al. 2017). In such cases, prediction performance based on variable selection methods like BL was preferred (Perez et al., 2012; Thavamanikumar, Dolferus and Thumma, 2015). As shown for Kernel Weight and Wheat Protein, in both same-year cross validations and cross-year predictions (Figure 1 and Figure 2) the advantage of BL for end-use quality traits was generally acknowledged. However, BL is much more computationally intensive than its more efficient competitors RF and RKHS that showed similar performance in this study.

Thanks to the advancement of genotyping capacity, evidence of biometric epistasis is growing (Costa et al., 2004; Mao et al., 2011; Hu et al., 2011; Muñoz et al., 2014); the role of physiological epistasis also is increasingly important (Boyle, Li and Pritchard, 2017). As in GS, the capacity to model the non-linear effects of all predictors would be beneficial (Howard et al., 2014; Gamal El-Dien et al., 2016). Also emphasized in Gianola et al. (2006), nonparametric algorithms like RKHS are advantageous when epistatic genetic architecture contributes significantly to phenotypic variation; similar results were supported by Battenfield et al. (2016), in traits like grain protein, grain hardiness and flour sedimentation phenotypes. On the contrary, except for the slight advantage in the cross-year prediction for Grain Yield (Figure 2), our results did not support the use of non-parametric methods like RKHS; even in cases of cross-validations where environmental variation is assumed to be constant, RKHS predictability was significantly outcompeted for the Grain Yield trait in 2016 (Figure 1). Abiotic stress, such as recurrent droughts in the southern Great Plains of the US, is a strong selective force. Given the inconsistent and insufficient rainfall patterns, selection for drought tolerance and avoidance would require strong directional selection to elevate allele frequency of stress tolerance genes (Böndel et al., 2018). However, breeding for individuals with drought tolerance might have depleted overall genetic diversity in regional breeding programs, and disrupted the genotype-phenotype (G-P) association for polygenic variation as a consequence (Assis et al., 2016; Jones et al., 2014; Pavlicev et al., 2010). Further, under directional selection, epigenetic gene action would be translated primarily into additive genetic variance of a small number of large fitness related QTLs (Crow 2010; Monnahan and Kelly, 2015), making RKHS a less preferential choice for GS for Grain Yield in a rapidly changing environment.

### Application of GS in Variety Development Program

Using a HRW wheat VDP as an example, breeding programs start out from the first generation of hybridization, and progenies then advance through three stages of inbreeding and population development, line development, and line testing, in which the genetic diversity within a VDP goes from highly heterogeneous, heterozygous populations in early generations to highly homozygous, homogeneous lines. Techniques like double haploid (DH) has been considered for rapidly developing inbred lines (Forster et al., 2007). Newly derived inbred lines are usually evaluated for 3 to 6 growing seasons before final selection to accomplish a specific goal or goals of the VDP.

Averaged across all three years of cross validation, GS predictability for Grain Yield was 0.52 (Figure 1); when evaluating GS performance across growing seasons, predictability decreased to an average of 0.25 (Figure 2). This over-inflated prediction accuracy was most obvious with Wheat Protein, and least so with SDS Sedimentation Volume (Figure 1 and Figure 2). Using a similar comparison, Battenfield et al. (2016) showed that cross-validation accuracy was overestimated by as much as 44% (1000-kernel weight) than forward prediction. Unlike Battenfield et al. (2016) that prediction accuracy was summarized from a number of full-sib families, whereas in our bi-parental DH population, the observed over-inflation would be more likely due to the inability to account for G×E, or genotype-by-environment interaction.

Accurate and stable selection of superior breeding lines over experimental trials is obviously important and desirable for VDP, but often difficult to achieve due to the presence of environmental variability. To ensure long-term response to selection, our results show that there are, however, cases where PS would be still preferential or cases that retraining with updated phenotypes should be performed. In principle, the superiority of GS was most notable when the selection intensity was high, and when large training information was available (Figure 3–5). In our case study when predicting line performance under sub-optimal conditions (*i*.*e*., drought conditions in this study) by information trained in normal growing conditions, additional phenotyping under the target, sub-optimal environment would be required to achieve a desirable selection response; this was most obvious for Grain Yield when selection intensity was high (Figure 3A). It is interesting to note that, as suggested by our results, training conducted in sub-optimal conditions could still provide GS predictability, while maintaining a desirable response to selection for both Grain Yield and SDS Sedimentation Volume (Figure 3–4). In another words, when making selection decisions for trials under unexpected environmental stress, like the frequent drought in the southern Great Plains of USA, using GS trained in optimal growing conditions could very likely result in unreliable outcomes. A similar deficiency of GS was also advised in Michel et al. (2018).

Further, effectiveness of GS has been examined with respect to the composition of training information (Rincent et al., 2012; Akdemir et al., 2015; Isidro et al., 2015; Michel et al., 2017a; Neyhart et al., 2017). A straightforward approach to optimize TP for prediction was to maximize phenotypic variation, as Isidro et al. (2015), Michel et al. (2017a), and Neyhart et al. (2017) all suggested an upward performance improvement in GS by the TP formed with two-tailed phenotypic data (*i*.*e*., highest and lowest phenotypes). In addition to this two-tailed TP design (GS_DTP_Ptails in Figure 3–6), our study also investigated various approaches of constructing DTP, such as two-tailed GEBVs (GS_DTP_Gtails in Figure 3–6) and the DTP formed by the majority votes of both GEBVs and raw phenotypes (GS_DTP_Htails in Figure 3–6). Using Grain Yield as an example for polygenic traits, a broadly appropriate guideline is, when training was obtained from normal growing conditions, straightforward GS approaches (GS_DTP_Gtails in Figure 3B) with an intermediate size of training population should be considered for high selection intensity; and when training was performed in a stressed growing condition, at a high selection intensity, optimized DTP with the majority votes could result in long-term advantage (GS_DTP_Htails, *m* = 40 and 60% in Figure 3A). The later scenario was as well beneficial for oligo-genic traits like SDS Sedimentation Volume (Figure 4A) and Kernel Weight (Figure 5A). Also with a heritability estimate of 0.74 (Table 2) and appreciable phenotypic correlation coefficients (Table 1), the stability in GS performance and in the response to selection across environment variability, makes SDS Sedimentation Volume a worthy candidate for GS in wheat variety development programs.

## Supplementary Materials

Supplementary Figure S1. Trace plots of residual variance from Bayesian Lasso for Grain Yield, SDS Sedimentation Volume, Kernel Weight, and Wheat Protein over three years.

Supplementary Figure S2. Trace plots of regularization parameter from Bayesian Lasso for Grain Yield, SDS Sedimentation Volume, Kernel Weight, and Wheat Protein over three years.

Supplementary Figure S3. Trace plots of residual variance from Bayesian reproducing kernel Hilbert space (Bayesian RKHS) for Grain Yield, SDS Sedimentation Volume, Kernel Weight, and Wheat Protein over three years.

Supplementary Figure S4. Trace plots of genetic variance from Bayesian reproducing kernel Hilbert space (Bayesian RKHS) for Grain Yield, SDS Sedimentation Volume, Kernel Weight, and Wheat Protein over three years.

Supplementary Figure S5. Average mean squared error of same-year cross validation for Grain Yield, SDS Sedimentation Volume, Kernel Weight, and Wheat Protein.

Supplementary Figure S6. Average mean squared error of cross-year prediction for Grain Yield, SDS Sedimentation Volume, Kernel Weight, and Wheat Protein. The year before and after dash stands for training and validation year respectively.

### Acknowledgements

Funding for this work was supported by grants from the Oklahoma Wheat Research Foundation (for X.H., R.F., C.P., L.Y., and C.C.), Oklahoma Center for the Advancement of Science and Technology (OCAST) award number PS15-011-2 for C.C.. This research represents the research outcomes for the USDA HATCH project OKL03011 (C.C.). Genotyping effort of this manuscript was supported from the National Science Foundation award number NSF-MRI 1626257 (C.C.).

## Conflict of Interest

The authors declare that they have no conflict of interest.

## Abbreviations

BL: Bayesian LASSO
CV: cross validation
DH: double haploid
DTP: designed training population
GBS: genotyping-by-sequencing
GEBV: genomic estimated breeding value
G x E: genotype-by-environment interaction
GS: genomic selection
GS_ATP: genomic selection by all training population
GS_DTP_Ptails: genomic selection by designed training population with phenotypic tails
GS_DTP_Gtails: genomic selection by designed training population with genomic tails
GS_DTP_Ptails: genomic selection by designed training population with hybrid tails
HRW: hard red winter
LD: linkage disequilibrium
kNN: k-nearest neighbor
MAF: minor allele frequency
MAS: marker assisted selection
MSE: mean squared error
NGS: next-generation sequencing
OAES: Oklahoma Agriculture Experiment Station
PCOR: Pearson’s correlation coefficient
PS: phenotypic selection
QTL: quantitative trait locus
RF: random forest
RKHS: reproducing kernel Hilbert space
RR: ridge regression
RRBLUP: ridge regression best linear unbiased prediction
SD: standard deviation
SNP: single-nucleotide polymorphism
TP: training population
VDP: variety development program
VP: validation population

